# Ribozyme-based biosensor for imaging gene expression in plants

**DOI:** 10.1101/2024.09.30.615876

**Authors:** Yang Liu, Ruchika Rajput, Torikul Islam, Ilenne Del Valle, Tao Yao, Rekha Agarwal, Brandon A. Boone, Carrie Eckert, Paul E. Abraham, Jin-Gui Chen, Gerald A. Tuskan, Xiaohan Yang

## Abstract

Detection of gene expression in plants is critical for understanding the molecular basis of complex plant biosystems and plant responses to environmental stresses. Here, we report the development of a split ribozyme-based biosensor that enables *in vivo* visualization of gene expression in plants. We demonstrated the utility of this biosensor in transient expression experiments (i.e., leaf infiltration in *Nicotiana benthamiana*) to detect RNAs derived from transgenes and tobacco rattle virus, respectively. Furthermore, we successfully engineered a split ribozyme-based biosensor in *Arabidopsis thaliana* for *in vivo* visualization of endogenous gene expression at the cellular level. In addition, we developed a platform for easy incorporation of different reporters into the RNA biosensor.

## Main

RNA plays a critical role in plant cellular activities and phenotypes by serving as messengers (i.e., mRNAs) to translate genetic information into proteins or as modulators (e.g., non-coding RNAs) to regulate gene expression ^1-3^. Technologies able to capture spatial-temporal dynamics of RNA molecules could be leveraged to unravel the molecular basis of complex phenotypes in plants or facilitate early detection of stressors affecting plants, developmental changes, and/or ectopic expression of transgenes. Current technologies for RNA analyses, such as quantitative reverse transcription polymerase chain reaction, *in situ* hybridization, and transcriptome-sequencing ^4-7^ require destructive, labor-intensive, and time-consuming measurements. Recent advancements in biosensors offer an alternative, non-destructive approach for measuring cellular or molecular activities ^8^. For example, RNA labeling technologies using aptamers have been applied for building biosensors for detection of gene expression ^9, 10^. However, these technologies require modification of input RNA signals and are thus not feasible for monitoring temporal and spatial patterns of endogenous gene expression in plants. Therefore, there is a pressing need for innovations in biosensors to enable *in-vivo* detection of RNA molecules within plants.

It was recently demonstrated that transcriptional signals could be monitored in mammalian and microbial cells using a synthetically split ribozyme-based biosensor that links native RNA signals to orthogonal protein outputs ^11-13^. These split ribozyme biosensors use a ribozyme derived from the *Tetrahymena* group I intron, which can catalyze trans-splicing of RNA molecules. To enable biosensing, the *Tetrahymena* ribozyme is split into two inactive fragments. The fragments only become functional when brought together by additional RNA interactions mediated by guide RNAs (gRNAs). The split ribozyme reassembly and subsequent splicing occur exclusively in the presence of a target RNA, ensuring specificity in detection. Moreover, because the system is linked to an orthogonal output, a single binding event between the target RNA and the split ribozyme reporter can initiate the translation of multiple copies of the reporter, acting as a signal amplifier. However, to our knowledge, the splicing activity of the ribozyme and split ribozyme-based RNA sensors have not been tested in plants. Here, we demonstrated the ribozyme-mediated transcript splicing functioned efficiently in *Nicotiana benthamiana* using Superfolder GFP (sfGFP) as a reporter. Furthermore, we successfully used split ribozyme-based biosensors to detect various types of transcripts in plants using both transient expression in *N. benthamiana* leaves and stable transformation in *Arabidopsis thaliana*. To allow for facile conversion of target RNA signals to protein outputs without the need to redesign the ribozyme, we connected the current split ribozyme platform with other protein outputs via the combination of ribozyme splicing with a 2A protein cleavage mechanism.

To evaluate ribozyme splicing activity in plants, we used a fluorescence-based splicing assay originally developed in *Escherichia coli* (*E. coli*) ^11^. Specifically, the self-splicing *Tetrahymena thermophila* ribozyme was inserted into the coding sequence (CDS) of sfGFP, interrupting the translation of the functional protein ^11^ (Fig. 1a). A functional sfGFP transcript can only be translated when the ribozyme excises itself from the surrounding exons and joins the split segments together ^11^ (Fig. 1a). We used the 35s promoter to drive expression of sfGFP inserted with *T. thermophila* ribozyme with and without an RNA stable structure hairpin 14 (HP14) ^14^ (Fig. 1b, c). Our results demonstrated that the ribozyme functions efficiently for splicing of the sfGFP transcript in *N. benthamiana* as visualized under a fluorescence microscope/stereoscope (Fig. 1d, e and Extended Data Fig. 1). We found that removing the HP14 hairpin from the RNA stable structure in the original biosensor used in *E. coli* ^11^ did not affect the sfGFP fluorescence output of the biosensor in *N. benthamiana* (Figs. 1d vs. 1e), indicating that this element was not necessary for the ribozyme function in plants. Western blot analysis confirmed that the sfGFP transcript fragments flanking the ribozyme were able to reassemble through ribozyme-mediated splicing and be translated into full-length sfGFP (Fig. 1f).

**Fig 1.**
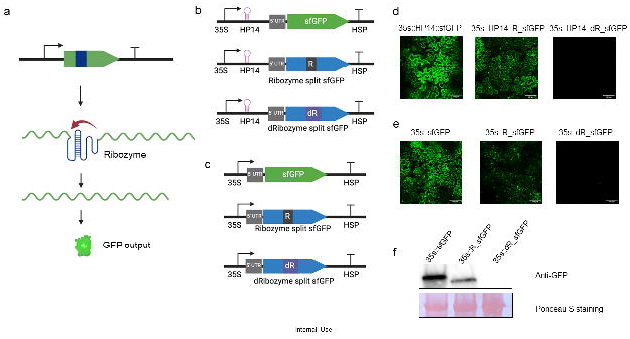
Evaluation of ribozyme splicing activity in *Nicotiana benthamiana* using sfGFP as the reporter via *Agrobacterium*-mediated leaf infiltration. **a**, Illustration of splicing activity assay using a sfGFP coding sequence (green) with the *Tetrahymena thermophila* ribozyme DNA sequence (dark blue) inserted within codon Y66. **b**, Construct design for testing ribozyme splicing activity in the presence of the HP14 hairpin structure using sfGFP as the reporter. 35S: Promoter region of the Cauliflower mosaic virus *35S* gene; 5’UTR: 5′ untranslated region of *Arabidopsis thaliana COLD REGULATED 47* gene; HSP: terminator region of *A. thaliana HEAT SHOCK PROTEIN 18*.*2* gene; R: ribozyme; dR: a catalytically dead G264A mutant ribozyme with IGS and PI loop removed; sfGFP: super folding *GREEN FLUORESCENT PROTEIN*. **c**, Construct design for testing ribozyme splicing activity without HP14 hairpin structure using sfGFP as the reporter. **d**, Evaluation of ribozyme splicing activity with sfGFP as reporter in the presence of HP14 in *N. benthamiana*. **e**, Evaluation of ribozyme splicing activity with sfGFP as reporter in the absence of HP14 in *N. benthamiana*. **f**, Western blot analysis of sfGFP resulting from the reassembly of sfGFP fragments caused by ribozyme-mediated transcript splicing.

Next, we aimed to convert RNA signals into orthogonal protein outputs in plants using the split ribozyme system. The core concept of the split ribozyme system is to attach guide RNA (gRNA) sequences to each ribozyme fragment for complementation. These gRNA sequences are engineered to base pair with a transcript and thereby, in the presence of the RNA target, the ribozyme fragments are brought together, enabling splicing to occur (Fig. 2a). We tested the split site at nucleotide 15 validated in *E. coli* ^11^ in *N. benthamiana* using *Agrobacterium*-mediated leaf infiltration. Specifically, we constructed two plasmid vectors (pXYB082, pXYB083), each carrying three expression cassettes. One cassette encodes the sequence of a red fluorescence protein (RFP), while the other two cassettes contain a fragment of sfGFP fused to a fragment of the *Tetrahymena* ribozyme linked to a gRNA targeting the RFP coding sequence (Fig. 2b). The two plasmid vectors are identical with the exception of gRNAs of differing lengths, 41-nt and 82-nt in pXYB082 and pXYB083, respectively. After leaf infiltration, green sfGFP fluorescence signal was detected in the *N. benthamiana* leaf tissue infiltrated with pXYB082 (containing a 41-nt gRNA) and pXYB083 (containing an 82-nt gRNA) (Fig. 2c and Extended Data Fig. 2a). Because low “off” state fluorescence from the split ribozyme system was consistently detected in *E. coli*, we would like to evaluate the potential background noise of split ribozyme system in plant using the plasmid vector (pXYB089) without any gRNAs. Weak sfGFP signal was also detected in the leaf tissue infiltrated with the negative control plasmid vector (pXYB089), but the GFP signal is significantly lower compared with split-ribozyme systems with gRNAs (Fig. 2c, d). This result indicates that the split ribozyme-based biosensor can be used to detect RNAs in plant cells.

**Fig 2.**
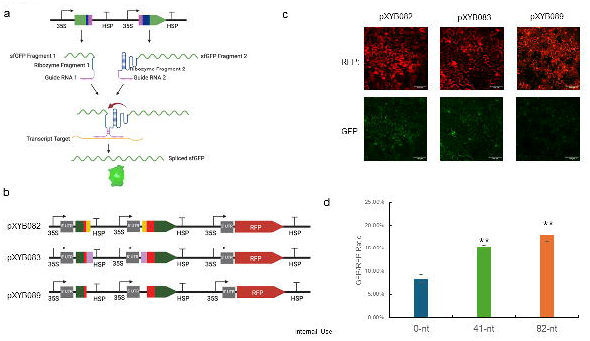
Split ribozyme-based biosensor system for RNA detection in plants using transient expression. **a**, Illustration of the split-ribozyme based biosensor system using a transcript input to produce functional sfGFP. **b**, Construct design for evaluation of the function of split ribozyme in plants using red fluorescent protein as transcript input. 35S: Promoter region of the Cauliflower mosaic virus *35S* gene; 5’UTR: 5′ untranslated region of *Arabidopsis thaliana COLD REGULATED 47* gene; HSP: terminator region of *Arabidopsis thaliana HEAT SHOCK PROTEIN 18*.*2* gene; RFP: *RED FLUORESCENT PROTEIN*. Green rectangles represent split sfGFP fragments, red rectangles represent split ribozyme fragments, yellow shapes represent two 41-nt gRNAs targeting RFP, pink rectangles represent two 82-nt gRNAs targeting RFP. **c**, Split ribozyme-based biosensor system for detecting RFP transcripts in *N. benthamiana* using sfGFP as reporter via *Agrobacterium*-mediated leaf infiltration. **d**, Characterization of fluorescence intensity for different lengths of guide RNAs targeting RFP. Error bars indicate error bars standard deviation of n□=□3 biological replicates.

To investigate if the increased gRNA length could enhance the florescent signal from the split-ribozyme system, we tested three more gRNAs with length of 123-nt (in pXYB084), 164-nt (in pXYB085), and 325-nt (in pXYB086), respectively. Surprisingly, unlike the results seen in *E. coli* ^11^, we did not observe an increase in the sfGFP fluorescence signal produced from split ribozyme system with increased gRNA length (Extended Data Fig. 2b).

We then tested the split ribozyme-based biosensor system targeting an *Arabidopsis thaliana* gene, *PLT1* (AT3G20840), which encodes an AP2 transcription factor expressed in the root quiescent center ^15^ using *N. benthamiana* leaf infiltration. We cloned the AT3G20840 coding sequence under a strong promoter 35s (pXYB151). We also created a biosensor vector with 82-nt gRNAs targeting the AT3G20840 coding sequence (pXYB147) (Fig. 3a). Green sfGFP signal was detected in the *N. benthamiana* leaf tissue co-infiltrated with pXYB147 and pXYB151 whereas sfGFP signal was not detected in the leaf tissue infiltrated with pXYB147 or pXYB151 alone (Fig. 3b and Extended Data Fig. 3a). Furthermore, we used the split ribozyme-based biosensor to detect infection by tobacco rattle virus (TRV). We engineered two biosensor vectors (pXYB155 and pXYB156) with 82-nt gRNAs targeting different regions of the TRV RNA sequence (Fig 3a). sfGFP signals were detected in the *N. benthamiana* leaf tissues co-infiltrated with TRV and one of the two biosensor constructs (pXYB155 and pXYB156). No sfGFP signal was detected in the leaf tissues infiltrated with TRV or the biosensor (pXYB155 or pXYB156) alone (Fig. 3c and Extended Data Fig. 3b). These results indicate that the split ribozyme-based biosensor system can be used *in vivo* to detect RNAs derived from diverse sources using transient expression approaches.

**Fig 3.**
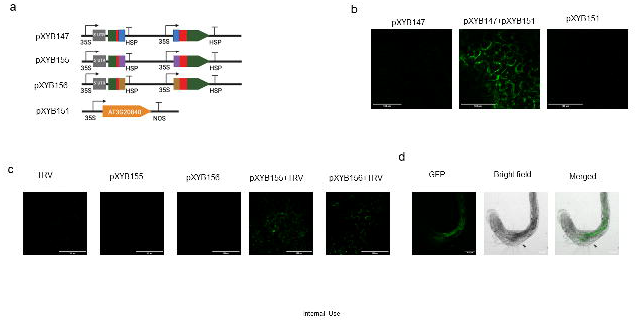
Utilizing the split ribozyme-based biosensor for imaging different RNA signals in plants. **a**, Construct design for detection of various types of RNA signals in plants. 35S: Promoter region of the Cauliflower mosaic virus *35S* gene; 5’UTR: 5′ untranslated region of *Arabidopsis thaliana COLD REGULATED 47* gene; HSP: terminator region of *Arabidopsis thaliana HEAT SHOCK PROTEIN 18*.*2* gene; blue rectangles represent 82-nt gRNAs targeting *Arabidopsis thaliana* gene *PLT1* (AT3G20840), and purple and brown rectangles represent 82-nt gRNAs targeting different regions of the tobacco rattle virus (TRV) RNA. **b**, Split ribozyme-based biosensor system for detecting the AT3G20840 transcript in *N. benthamiana* using sfGFP as reporter via *Agrobacterium*-mediated leaf infiltration. **c**, Testing the split ribozyme-based biosensor system for detecting the TRV RNA in *N. benthamiana* using sfGFP as reporter via *Agrobacterium*-mediated leaf infiltration. **d**, Split ribozyme-based biosensor system for detecting the expression of an endogenous gene (i.e., AT3G20840) in the roots of transgenic *A. thaliana* plants engineered with the construct pXYB147 using *Agrobacterium*-mediated stable transformation.

Based on the success of utilizing the split ribozyme-based biosensor system to detect RNAs through transient expression in *N. benthamiana* leaves, we engineered the biosensor construct pXYB147 (Fig. 3a) with 82-nt gRNAs targeting the AT3G20840 coding sequence into *A. thaliana* using *Agrobacterium*-mediated stable transformation. The sfGFP signal was detected in the roots of three T1 transgenic *Arabidopsis* plants (Fig. 3d and Extended Data Fig. 4), indicating that the split ribozyme-based biosensor system can be used to detect endogenous gene expression at the cellular level in plants with high specificity.

**Fig 4.**
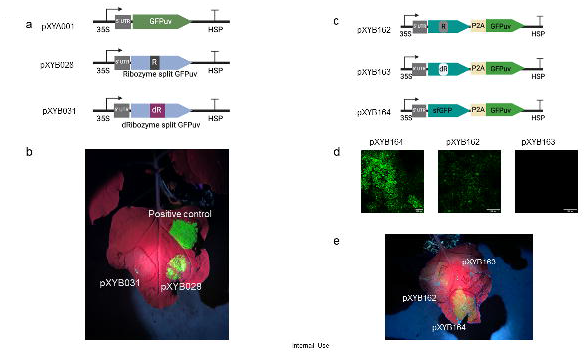
Engineering the ribozyme system for interchangeable protein outputs in *Nicotiana benthamiana* using GFPuv as the reporter via *Agrobacterium*-mediated leaf infiltration. **a**, Construct design for testing the reassembly of GFPuv fragments mediated by ribozyme splicing. GFPuv: enhanced *YELLOW GREEN FLUORESCENT LIKE PROTEIN* that can be detected by unaided eyes under ultraviolet light. **b**, Evidence of the reassembly of GFPuv fragments mediated by ribozyme splicing in *N. benthamiana*. **c**, Construct design for testing the ribozyme splicing activity with sfGFP-P2A-GFPuv fusion protein as the reporter. P2A: a 2A self-cleaving peptide; 35S: Promoter region of the Cauliflower mosaic virus *35S* gene; 5’UTR: 5′ untranslated region of *Arabidopsis thaliana COLD REGULATED 47* gene; HSP: terminator region of *Arabidopsis thaliana HEAT SHOCK PROTEIN 18*.*2* gene; R: Ribozyme; dR: a catalytically dead G264A mutant ribozyme with IGS and PI loop removed. **d**, Ribozyme-mediated reassembly of sfGFP transcript connected to eYGFP via P2A. **e**, Visualization of eYGFPuv, connected via P2A, to sfGFP reassembled via ribozyme splicing in *N. benthamiana*.

RNA detection using the split ribozyme-based biosensor system with sfGFP as a reporter requires the use of a fluorescence microscope, which is technically demanding and time consuming and does not allow detection in live tissues. To address this limitation, we replaced sfGFP with a GFP-like protein (eYGFPuv), which was recently utilized for *in-vivo* visualization of transgene expression under a UV flashlight ^16^. First, we tested if ribozyme-mediated splicing can reassemble two partial fragments of the eYGFPuv transcripts in plant cells using *N. benthamiana* leaf infiltration. Ribozyme-mediated cleavage at the 5′ splice site depends on formation of a base-paired helix P1 between the Internal Guide Sequence (IGS) and sequences adjacent the splice site, and the presence of a U·G “wobble” base-pair positioned 4-6 residues from the base of this helix defines the susceptible phosphodiester bond ^17^. Therefore, the ribozyme must be inserted immediately downstream of an uracil where the IGS should start with a guanine and the remaining IGS sequence is in reverse complementation to the first 5 base pairs of the P1 helix ^17^. Following these principles, we tested different insertion sites of eYGFPuv coding sequence and designed alternate IGS for different target sites. We discovered that the eYGFPuv with the ribozyme inserted after the last amino acid of Y72 showed a strong fluorescence signal under a UV flashlight (Fig. 4a, b). However, only one of six replicates showed desirable eYGFPuv signal, indicating correct splicing efficiency is low. To enhance the ribozyme splicing efficiency, we modified the IGS of the original ribozyme design, known to pair with the P1 helix, and tested different RNA stable motif in eukaryotes (Extended Data Fig.5). However, none of those optimizations efficiently enhanced the ribozyme splicing activity in eYGFPuv (Extended Data Fig.5). These results suggest that ribozyme splicing efficiency is different under different transcript context in plants. Variable ribozyme-mediated splicing efficiency was also observed in mammalian cells ^18, 19^. Hence, a modular platform allowing for simple exchange of the output protein would be helpful for avoiding the inconsistent ribozyme splicing activity in different context.

To bypass the need to redesign ribozyme insertion sites for different protein outputs (e.g., eYGFPuv), we created a construct (pXYB162) with eYGFPuv attached via the 2A self-cleavage peptide P2A to sfGFP (Fig. 4c), which was optimized for ribozyme-mediated splicing. The sfGFP signal was detected in the *N. benthamiana* leaf tissue infiltrated with the construct pXYB162 (containing two sfGFP fragments flanking the ribozyme) although the signal was weaker than that in leaf tissue infiltrated with the positive control construct pXYB164 (containing intact sfGFP); no sfGFP signal was detected in the leaf tissue infiltrated with the negative control construct pXYB163 (containing two sfGFP fragments flanking a catalytically dead G264A mutant ribozyme with IGS and PI loop removed) (Fig. 4d and Extended Data Fig. 6). This result indicates that the eYGFPuv attachment did not affect the reassembly of sfGFP fragments through ribozyme-mediated transcript splicing.

While the eYGFPuv fluorescence signal detected in *N. benthamiana* leaf tissue infiltrated with pXYB162 was much weaker than that in leaf tissue infiltrated with the positive control construct pXYB164, we do demonstrate that ribozyme-mediate splicing and 2A self-cleavage peptide can be combined and still produce functional proteins (Fig. 4e). Therefore, the sfGFP-2A-protein’s configuration could potentially be used for designing a new split ribozyme-based system with any functional protein as an output for RNA-based genome engineering at cellular level in plants, although further work is necessary to optimize the sfGFP-2A-protein configuration for optimizing expression of the second functional protein after 2A splicing.

In conclusion, we have successfully demonstrated that the split ribozyme-based biosensor system can be used for detecting RNAs in plants through both transient expression and stable transformation. We call this biosystem “Plant RNA Vision”, which is expected to have broad applications in plant biology and plant biotechnology, such as facilitating the experimental validation of tissue/cell-type specific gene expression predicted by single-cell transcriptome-sequencing ^7^ and monitoring plant gene expression responses to abiotic and biotic factors (Extended Data Fig. 7). Furthermore, the Plant RNA Vision system holds great potential for building RNA-mediated tunable genetic circuits to establish new capabilities for plant synthetic biology research.

## Methods

### Plant materials

*Arabidopsis* wild-type Col-0 plants and *Nicotiana benthamiana* (GenBank: PRJNA170566) plants were grown in the soil in the growth chamber at 21°C under 100 μmol m^−2^ s^−1^ white light with 12-h light/12-h dark photoperiod.

### Plasmids construction

All the DNA constructs used in this study are listed in Supplementary Table 1, which were created via Gibson assembly using NEBuilder^®^ HiFi DNA Assembly master mix (NEB, Cat. No. E2621) or Golden Gate assembly using the Modular Vector Library cloning Kit from Dr. Dan Voytas’ lab at the University of Minnesota ^20^. Sequence information for the DNA constructs was provided in Supplementary Table 2 to 5. Guide RNA sequences design was illustrated in Extended Data Fig. 8. Sequences were synthesized by TWIST Bioscience (South San Francisco, United States).**Tobacco leaf infiltration**

The *Agrobacterium tumefaciens* strain GV3101 harboring the plasmid of interest was infiltrated into the leaves of five-to six-week-old *N. benthamiana* plants using a 1mL syringe without a needle as described previously ^21, 22^. Briefly, cultures containing the vector of interest were grown in lysogeny broth (LB) medium for 36 - 48 hours and resuspended in agroinfiltration buffer (10□mM MES, pH 5.6, 10□mM MgCl2 and 200□µM acetosyringone) to an optical density of 0.5 at 600□nm (OD600). Bacterial suspensions were incubated for 2 to 4 hours at room temperature and then mixed for co-expression experiments. Agroinfiltrations were carried out through the abaxial surface of the three youngest fully expanded leaves of each plant with a 1-ml needle-free syringe. At least two independent experiments were conducted for each treatment. Each treatment includes six replicates.

### Stable transformation in *Arabidopsis thaliana*

The *Agrobacterium* strain ‘GV3101’ was used for the transformation of *A. thaliana* wild type ‘Col-0’ via the floral dip method with modification as described previously ^16^. T1 seeds co-transformed with two constructs were selected on K1 medium (1L 2.154 g MS, 100 mg myo-insoitol, 0.5g MES, 10 g sucrose, 8 g agar if plates; based on the recommendation of ABRC (Arabidopsis Biological Resource Center) (https://abrc.osu.edu/seed-handling) for Arabidopsis seed germination) with 50 mg/L Kanamycin and 25 mg/L Hygromycin. Seeds were put under dark at 4°C for 2 days and then germinated for 6 to 8 days under 100 to 150 µmol m^−2^ s^−1^ fluorescent warm white light, 12h light/12h dark, 20°C, and 70% humidity. Then, seedlings were collected for GFP visualization.

### Fluorescence measurements

The fluorescence of sfGFP and RFP was visualized and imaged using a Zeiss LSM 710 confocal microscope and images were analyzed with the images were analyzed with Fiji software ^23^. GFPuv signal was visualized using LIGHTFE UV302D (365□nm) ^16^.

### Statistical analysis

The sample average and standard deviation (s.d.) were calculated from three replicates of each treatment. Significance in all experiments was assessed using a T-Test with an alpha of 0.05. In all cases, resulting p-values of less than 0.05 for fluorescence intensity of experimental treatment compared to negative control were considered significantly different.

### Western blot

Leaf discs from infiltrated tobacco leaves were harvested and ground into a fine powder in liquid nitrogen. Total proteins were then extracted by adding 2x Laemmli sample buffer (BioRad) and boiling for 10 minutes. Cell debris was removed by centrifugation at 16,000 g for 10 minutes at 4°C. The supernatant was resolved on a 4-20% SDS–PAGE gel (BioRad) and transferred to polyvinylidene fluoride membranes (Millipore). Anti-GFP polyclonal antibodies (Invitrogen) were used as primary antibodies at a 1:2,000 (v/v) dilution. Immunoblots were detected using peroxidase-conjugated anti-rabbit IgG (Sigma-Aldrich) at a 1:7,500 (v/v) dilution and ECL substrate (Thermo Scientific) with the BioRad ChemiDoc gel documentation system.

## Data availability

The plasmids generated in this study will be available at Addgene (https://www.addgene.org/).

## Acknowledgements

The writing of this manuscript was supported by the U.S. Department of Energy (DOE) Genomic Science Program, as part of the Secure Ecosystem Engineering and Design (SEED) Scientific Focus Area. This material is also based upon work supported by the Center for Bioenergy Innovation (CBI), U.S. Department of Energy, Office of Science, Biological and Environmental Research Program under Award Number ERKP886. Oak Ridge National Laboratory is managed by UT-Battelle, LLC for the U.S. Department of Energy under Contract Number DE-AC05-00OR22725.

## Author contributions

X.Y. conceived, designed, and supervised the project, and reviewed and edited the manuscript. G.A.T. supervised the project and reviewed and edited the manuscript. Y.L. conceived, designed, and performed experiments, analyzed the data, and wrote the manuscript. R.R., T.I., and R.A. performed the experiments in Arabidopsis transformation. T.Y. performed the western blot experiments. B.A.B. helped with confocal data analysis. I.D.V. and C.A.E. provided information for initiating the project and offered counseling service for some of the experiments. All authors read and contributed to the content, edited, or reviewed it.

