## Supplementary information for "Ribozyme-based biosensor for imaging gene expression in plants"


| Tables | Title | Page |
|---|---|---|
| 1 | All plasmids used in this study | 1 - 3 |
| 2 | Example Plant RNA Vision plasmid sequence | 3 - 5 |
| 3 | Transcripts inputs used in this study | 5 - 7 |
| 4 | Guide RNA sequences used in this study | 7 - 8 |
| 5 | Sequences of other biological components used in this study | 8 - 9 |


**Supplementary Table 1. List of plasmids used in this study.** 35S: Promoter region of the Cauliflower mosaic virus *35S* gene, 5'UTR: 5′ untranslated region of *Arabidopsis thaliana COLD REGULATED 47* gene system, HSP: terminator region of *Arabidopsis thaliana HEAT SHOCK PROTEIN 18.2* gene, NOS: terminator region of *Agrobacterium NOPALINE SYNTHASE* gene, RFP: *RED FLUORESCENT PROTEIN*. Green shapes represent split sfGFP fragments, red shapes represent split ribozyme fragments, yellow shapes represent two gRNAs with length at 41bp targeting RFP, pink shapes represent two gRNAs with length at 82bp targeting RFP, blue shapes represent gRNAs with length at 82bp targeting *Arabidopsis thaliana* gene *AT3G20840*.

| Plasmid | Figures | Description | Plasmid design |
|---|---|---|---|
| pXYB033 | Fig. 1 | sfGFP expression | 35S::5'UTR::sfGFP::HSP |
| pXYB034 | Fig. 1 | Ribozyme split sfGFP expression | 35S::5'UTR::sfGFPn_Y66-Ribozyme-2nd_sfGFPc::HSP |
| pXYB035 | Fig. 1 | Deactivated ribozyme split sfGFP expression | 35S::5'UTR::sfGFPn_Y66-dRibozyme-2nd_sfGFPc::HSP |
| pAXY0001 | Fig. 4 | GFPuv expression | 35S::5'UTR::GFPuv::HSP |
| pXYB028 | Fig. 4 | Ribozyme split GFPuv expression | 35S::5'UTR::GFPuvn_Y72-Ribozyme-2nd_GFPuvc::HSP |
| pXYB031 | Fig. 4 | Deactivated ribozyme split GFPuv expression | 35S::5'UTR::GFPuvn_Y72-dRibozyme-2nd_GFPuvc::HSP |
| pXYB082 | Fig. 2 | Plant RNA Vision constructs with 41 bp gRNAs together with RFP input | 35S::5'UTR::sfGFPn_Y66-P1_loop_01-stem01-gRNA01::HSP --- 35S::5'UTR::gRNA02-stem_02-P1_loop_03-internal guide sequence-Ribozyme-sfGFPn_Y66::HSP--- 35S::RFP::HSP |
| pXYB083 | Fig. 2 | Plant RNA Vision constructs with 82 bp gRNAs together with RFP input | 35S::5'UTR::sfGFPn_Y66-P1_loop_01-stem01-gRNA03::HSP --- 35S::5'UTR::gRNA04-stem_02-P1_loop_03-internal guide sequence-Ribozyme-sfGFPn_Y66::HSP--- 35S::RFP::HSP |
| pXYB089 | Fig. 2 | Plant RNA Vision constructs with no gRNAs together with RFP input | 35S::5'UTR::sfGFPn_Y66-P1_loop_01-stem01::HSP --- 35S::5'UTR::stem_02-P1_loop_03-internal guide sequence-Ribozyme-sfGFPn_Y66::HSP--- 35S::RFP::HSP |

| | | | |
|---|---|---|---|
| pXYB147 | Fig. 3 | Plant RNA Vision constructs with with 82 bp gRNAs targeting *AT3G20840* | 35S::5'UTR::sfGFPn_Y66-P1_loop_01-stem01-gRNA05::HSP --- 35S::gRNA06-stem_02-P1_loop_03-internal guide sequence-Ribozyme-sfGFPn_Y66::HSP |
| pXYB151 | Fig. 3 | Constitutive RNA input | 35S::*AT3G20840*::NOS |
| pXYB155 | Fig. 3 | Plant RNA Vision constructs with with 82 bp gRNAs targeting TRV | 35S::5'UTR::sfGFPn_Y66-P1_loop_01-stem01-gRNA13::HSP --- 35S::gRNA14-stem_02-P1_loop_03-internal guide sequence-Ribozyme-sfGFPn_Y66::HSP |
| pXYB156 | Fig. 3 | Plant RNA Vision constructs with with 82 bp gRNAs targeting TRV | 35S::5'UTR::sfGFPn_Y66-P1_loop_01-stem01-gRNA15::HSP --- 35S::gRNA16-stem_02-P1_loop_03-internal guide sequence-Ribozyme-sfGFPn_Y66::HSP |
| pXYB162 | Fig. 4 | Ribozyme split sfGFP linked with GFPuv via T2A expression | 35S::5'UTR::sfGFPn_Y66-Ribozyme-2nd_sfGFPc-T2A-GFPuv::HSP |
| pXYB163 | Fig. 4 | Deactivated ribozyme split sfGFP linked with GFPuv via T2A expression | 35S::5'UTR::sfGFPn_Y66-dRibozyme-2nd_sfGFPc-T2A-GFPuv::HSP |
| pXYB164 | Fig. 4 | sfGFP linked with GFPuv via T2A expression | 35S::5'UTR::sfGFP-T2A-GFPuv::HSP |
| pXYB084 | Extended Data Fig. 2 | Plant RNA Vision constructs with 123 bp gRNAs together with RFP input | 35S::5'UTR::sfGFPn_Y66-P1_loop_01-stem01-gRNA07::HSP --- 35S::5'UTR::gRNA08-stem_02-P1_loop_03-internal guide sequence-Ribozyme-sfGFPn_Y66::HSP---35S::RFP::HSP |
| pXYB085 | Extended Data Fig. 2 | Plant RNA Vision constructs with 164 bp gRNAs together with RFP input | 35S::5'UTR::sfGFPn_Y66-P1_loop_01-stem01-gRNA09::HSP --- 35S::5'UTR::gRNA10-stem_02-P1_loop_03-internal guide sequence-Ribozyme-sfGFPn_Y66::HSP---35S::RFP::HSP |
| pXYB086 | Extended Data Fig. 2 | Plant RNA Vision constructs with 325 bp gRNAs together with RFP input | 35S::5'UTR::sfGFPn_Y66-P1_loop_01-stem01-gRNA11::HSP --- 35S::5'UTR::gRNA12-stem_02-P1_loop_03-internal guide sequence-Ribozyme-sfGFPn_Y66::HSP---35S::RFP::HSP |
| pXYB105 | Extended Data Fig. 5 | GFPuv with 3' tEvopreq motif expression | 35S::5'UTR::GFPuv+tEvopreq::HSP |

| | | | |
|---|---|---|---|
| pXYB106 | Extended Data Fig. 5 | Ribozyme split GFPuv with tEvopreq motif expression | 35S::5'UTR::GFPuvn_Y72-Ribozyme-2nd_GFPuvc+tEvopreq::HSP |
| pXYB107 | Extended Data Fig. 5 | Deactivated ribozyme split GFPuv tEvopreq motif expression | 35S::5'UTR::GFPuvn_Y72-dRibozyme-2nd_GFPuvc+tEvopreq::HSP |
| pXYB108 | Extended Data Fig. 5 | GFPuv with 3' gq2 motif expression | 35S::5'UTR::GFPuv+gq2::HSP |
| pXYB109 | Extended Data Fig. 5 | Ribozyme split GFPuv with gq2 motif expression | 35S::5'UTR::GFPuvn_Y72-Ribozyme-2nd_GFPuvc+gq2::HSP |
| pXYB110 | Extended Data Fig. 5 | Deactivated ribozyme split sfGFP gq2 motif expression | 35S::5'UTR::GFPuvn_Y72-dRibozyme-2nd_GFPuvc+gq2::HSP |
| pXYB111 | Extended Data Fig. 5 | GFPuv with 3' Triplex motif expression | 35S::5'UTR::GFPuv+Triplex::HSP |
| pXYB112 | Extended Data Fig. 5 | Ribozyme split GFPuv with Triplex motif expression | 35S::5'UTR::GFPuvn_Y72-Ribozyme-2nd_GFPuvc+Triplex::HSP |
| pXYB113 | Extended Data Fig. 2 | Deactivated ribozyme split GFPuv with Triplex motif expression | 35S::5'UTR::GFPuvn_Y72-dRibozyme-2nd_GFPuvc+Triplex::HSP |
| pXYB176 | Extended Data Fig. 5 | Ribozyme with IGS2 split GFPuv expression | 35S::5'UTR::GFPuvn_Y72-Ribozyme-2nd_GFPuvc+Triplex::HSP |
| pXYB177 | Extended Data Fig. 5 | Deactivated ribozyme with IGS2 split GFPuv expression | 35S::5'UTR::GFPuvn_Y72-dRibozyme-2nd_GFPuvc+Triplex::HSP |

**Supplementary Table 2. Sequence of a representative DNA construct in the split ribozyme-based RNA biosensor system.** See construct design in Fig. 2b.

| Construct name | DNA sequence |
|---|---|
| pXYB147 (PlantRNA Vision construct with with 82 bp gRNAs targeting *Arabidopsis thaliana* gene *AT3G20840*) [35S::5'UTR::sfGFPn _Y66-P1_loop_01-stem01-gRNA05::HSP --- 35S::gRNA06-stem_02-P1_loop_02-internal guide | TGAGACTTTTCAACAAAGGGTAATATCCGGAAACCTCCTCGGATTC CATTGCCCAGCTATCTGTCACTTTATTGTGAAGATAGTGGAAAAGGA AGGTGGCTCCTACAAATGCCATCATTGCGATAAAGGAAAGGCCATC GTTGAAGATGCCTCTGCCGACAGTGGTCCCAAAGATGGACCCCCAC CCACGAGGAGCATCGTGGAAAAAGAAGACGTTCCAACCACGTCTT CAAAGCAAGTGGATTGATGTGATATCTCCACTGACGTAAGGGATGA CGCACAATCCCACTATCCTTCGCAAGACCCTTCCTCTATATAAGGAA GTTCATTTCATTTGGAGAGAACACGGGGGACTCAAACATTACTCAT TCACAAAACCATCTTAAAGCAACTACACAAGTCTTGAAATTTTCTC ATATTTTCTATTTACTATATAAACTTTTAATCAAATCAAGATTAAAGTT AATTAAATGAGCAAAGGAGAAGAACTTTTCACTGGAGTTGTCCCAA TTCTTGTTGAATTAGATGGTGATGTTAATGGGCACAAATTTTCTGTC CGTGGAGAGGGTGAAGGTGATGCTACAAACGGAAAACTCACCCTT |

| | |
|---|---|
| sequence-Ribozyme-sfGFPn_Y66::HSP] | AAATTTATTTGCACTACTGGAAAACTACCTGTTCCGTGGCCAACACT<br>TGTCACTACTCTGACCTAAATAGCAAGGATCACCGGTTTGCTCACA<br>CCGAGAAAATCGGCCACTTTTGGAACCTCTCCTCCTTCATCTCCTCC<br>TCCACCGTGTGGATTGATCATATATGAAGATGAAGATGAAATATTTG<br>GTGTGTCAAATAAAAAGCTTGTGTGCTTAAGTTTGTGTTTTTTTCTT<br>GGCTTGTTGTGTTATGAATTTGTGGCTTTTTCTAATATTAAATGAATG<br>TAAGATCTCATTATAATGAATAAACAAATGTTTCTATAATCCATTGTG<br>AATGTTTTGTTGGATCTCTTCTGCAGCATATAACTACTGTATGTGCTA<br>TGGTATGGACTATGGAATATGATTAAAGATAAGATGGGCTCATAGAG<br>TAAAACGAGGCGAGGGACCTATAAACCTCCCTTCATCATGCTATTTC<br>ATGATCTATTTTATAAAATAAAGATGTAGAAAAAAGTAAGCGTAATA<br>ACCGCAAAACAAATGATTTAAAACATGGCACATAATGAGGAGATTA<br>AGTTCGGTTTACGTTTATTTTAGTACTAATTGTAACGTGAGACTACG<br>TATCGGGAATCGCCTAATTAAAGCATTAATGCGAACCTGATTAGATT<br>CACCGACCCTCCTATCGTGTCGACCTTTCTGTTTCTTAGAATTTTTTG<br>GTAGTCTATGTACTAATAATGTCAGCTTCGTATTTATTTCATAAGCAA<br>TTTGCATTTGCAATTTGTTTTTTACTTTTATTTTTATTGTATTGTGGAA<br>TGTGGACTCGTACCAACATGAAGTTATATACCACCAAAAAAATTAC<br>AGTTAGTCAAAAGATTCACGAGTGAGAGCTACTTATGATTGTCTTTT<br>ACGTATATGTCTAATTGTCTATTTGCTCAATAATCTTTGTACTTTCTTT<br>TGTCGTTGATAAAATCACAAAGTTCCAAAAGTAATCGAATGATTTGC<br>TTTTAAGAAAAGAAGAGCTCAATAATTCAACATATATCTGTACACAG<br>ACGGACTGAGACTTTTCAACAAAGGGTAATATCCGGAAACCTCCTC<br>GGATTCCATTGCCCAGCTATCTGTCACTTTATTGTGAAGATAGTGGA<br>AAAGGAAGGTGGCTCCTACAAATGCCATCATTGCGATAAAGGAAAG<br>GCCATCGTTGAAGATGCCTCTGCCGACAGTGGTCCCAAAGATGGAC<br>CCCCACCCACGAGGAGCATCGTGGAAAAAGAAGACGTTCCAACCA<br>CGTCTTCAAAGCAAGTGGATTGATGTGATATCTCCACTGACGTAAG<br>GGATGACGCACAATCCCACTATCCTTCGCAAGACCCTTCCTCTATAT<br>AAGGAAGTTCATTTCATTTGGAGAGAACATTGATTGGACGCTAGGC<br>ATCAAGCTATTGGTATGGAAGTAGTAGTCTGAGTCGTTGTAAGCTAC<br>TAGGTGGTTGGATTGGTTTGATCCTAGTTACCTTTGGGTCAAAAAGT<br>TATCAGGCATGCACCTGGTAGCTAGTCTTTAAACCAATAGATTGCAT<br>CGGTTTAAAAGGCAAGACCGTCAAATTGCGGGAAAGGGGTCAACA<br>GCCGTTCAGTACCAAGTCTCAGGGGAAACTTTGAGATGGCCTTGCA<br>AAGGGTATGGTAATAAGCTGACGGACATGGTCCTAACCACGCAGCC<br>AAGTCCTAAGTCAACAGATCTTCTGTTGATATGGATGCAGTTCACAG<br>ACTAAATGTCGGTCGGGGAAGATGTATTCTTCTCATAAGATATAGTC<br>GGACCTCTCCTTAATGGGAGCTAGCGGATGAAGTGATGCAACACTG<br>GAGCCGCTGGGAACTAATTTGTATGCGAAAGTATATTGATTAGTTTT<br>GGAGTACTCGATGGTGTTCAATGCTTTTCCCGTTATCCGGATCACAT<br>GAAACGGCATGACTTTTTCAAGAGTGCCATGCCCGAAGGTTATGTA<br>CAGGAACGCACTATATCTTTCAAAGATGACGGGACCTACAAGACGC<br>GTGCTGAAGTCAAGTTTGAAGGTGATACCCTTGTTAATCGTATCGAG<br>TTAAAGGGTATTGATTTTAAAGAAGATGGAAACATTCTTGGACACA<br>AACTCGAGTACAACTTTAACTCACACAATGTATACATCACGGCAGA<br>CAAACAAAAGAATGGAATCAAAGCTAACTTCAAAATTCGCCACAA<br>CGTTGAAGATGGTTCCGTTCAACTAGCAGACCATTATCAACAAAAT<br>ACTCCAATTGGCGATGGCCCTGTCCTTTTACCAGACAACCATTACCT<br>GTCGACACAATCTGTCCTTTCGAAAGATCCCAACGAAAAGCGTGAC<br>CACATGGTCCTTCTTGAGTTTGTAACTGCTGCTGGGATTACACATGG |

| | | |
|---|---|---|
| | | CATGGATGAGCTCTACAAATAAATATGAAGATGAAGATGAAATATTT<br>GGTGTGTCAAATAAAAAGCTTGTGTGCTTAAGTTTGTGTTTTTTTCT<br>TGGCTTGTTGTGTTATGAATTTGTGGCTTTTTCTAATATTAAATGAAT<br>GTAAGATCTCATTATAATGAATAAACAAATGTTTCTATAATCCATTGT<br>GAATGTTTTGTTGGATCTCTTCTGCAGCATATAACTACTGTATGTGCT<br>ATGGTATGGACTATGGAATATGATTAAAGATAAGATGGGCTCATAGA<br>GTAAAACGAGGCGAGGGACCTATAAACCTCCCTTCATCATGCTATTT<br>CATGATCTATTTTATAAAATAAAGATGTAGAAAAAAGTAAGCGTAAT<br>AACCGCAAAACAAATGATTTAAAACATGGCACATAATGAGGAGATT<br>AAGTTCGGTTTACGTTTATTTTAGTACTAATTGTAACGTGAGACTAC<br>GTATCGGGAATCGCCTAATTAAAGCATTAATGCGAACCTGATTAGAT<br>TCACCGACCCTCCTATCGTGTCGACCTTTCTGTTTCTTAGAATTTTTT<br>GGTAGTCTATGTACTAATAATGTCAGCTTCGTATTTATTTCATAAGCA<br>ATTTGCATTTGCAATTTGTTTTTTACTTTTATTTTTATTGTATTGTGGA<br>ATGTGGACTCGTACCAACATGAAGTTATATACCACCAAAAAAATTAC<br>AGTTAGTCAAAAGATTCACGAGTGAGAGCTACTTATGATTGTCTTTT<br>ACGTATATGTCTAATTGTCTATTTGCTCAATAATCTTTGTACTTTCTTT<br>TGTCGTTGATAAAATCACAAAGTTCCAAAAGTAATCGAATGATTTGC<br>TTTTAAGAAAAGAAGAGCTCAATAATTCAACATATATCTGTACACA |

**Supplementary Table 3. Sequences of genes encoding the RNA targets used for testing the split ribozyme-based RNA biosensor system in plants. TRV represent Tabacco Rattle Virus.**

| Name | Length (nt) | Sequence |
|---|---|---|
| RFP | 678 | ATGGCGAGTAGCGAAGACGTTATCAAAGAGTTCATGCGTTTC<br>AAAGTTCGTATGGAAGGTTCCGTTAACGGTCACGAGTTCGAA<br>ATCGAAGGTGAAGGTGAAGGTCGTCCGTACGAAGGTACCCA<br>GACCGCTAAACTGAAAGTTACCAAAGGTGGTCCGCTGCCGT<br>TCGCTTGGGACATCCTGTCCCCGCAGTTCCAGTACGGTTCCA<br>AAGCTTACGTTAAACACCCGGCTGACATCCCGGACTACCTGA<br>AACTGTCCTTCCCGGAAGGTTTCAAATGGGAACGTGTTATGA<br>ACTTCGAAGACGGTGGTGTTGTTACCGTTACCCAGGACTCCT<br>CCCTGCAAGACGGTGAGTTCATCTACAAAGTTAAACTGCGTG<br>GTACCAACTTCCCGTCCGACGGTCCGGTTATGCAGAAAAAA<br>ACCATGGGTTGGGAAGCTTCCACCGAACGTATGTACCCGGA<br>AGACGGTGCTCTGAAAGGTGAAATCAAAATGCGTCTGAAAC<br>TGAAAGACGGTGGTCACTACGACGCTGAAGTTAAAACCACC<br>TACATGGCTAAAAAACCGGTTCAGCTGCCGGGTGCTTACAAA<br>ACCGACATCAAACTGGACATCACCTCCCACAACGAAGACTA<br>CACCATCGTTGAACAGTACGAACGTGCTGAAGGTCGTCACT<br>CCACCGGTGCTTAA |
| AT3G20840 | 2048 | GAGCTTGGTATATTGATCATACCAAAGTGGTAGTGATTTATTG<br>ATTAACCCAAACACAAAATAAACAGATTTGACTCAAAAAGA<br>AGAAAATGAATTCTAACAACTGGCTTGGCTTTCCTCTTTCAC<br>CGAACAACTCTTCTTTGCCTCCTCATGAATACAACCTTGGCTT<br>GGTCAGCGACCATATGGACAACCCTTTTCAAACACAAGAGT<br>GGAATATGATCAATCCACACGGTGGAGGAGGAGATGAAGGA<br>GGAGAGGTTCCAAAAGTGGCCGATTTTCTCGGTGTGAGCAA<br>ACCGGACGAAAACCAATCCAACCACCTAGTAGCTTACAACG<br>ACTCAGACTACTACTTCCATACCAATAGCTTGATGCCTAGCGT |

| | | |
|---|---|---|
| | | CCAATCAAACGATGTCGTTGTAGCAGCTTGTGACTCCAATAC<br>TCCTAACAACAGTAGCTATCATGAGCTTCAAGAGAGTGCTCA<br>CAATCTACAGTCACTTACTTTGTCCATGGGGACCACCGCTGG<br>TAATAATGTTGTAGACAAAGCTTCACCATCCGAGACCACCGG<br>GGATAACGCTAGCGGTGGAGCACTAGCCGTTGTTGAGACGG<br>CCACGCCAAGACGTGCATTGGACACTTTCGGACAACGAACC<br>TCGATCTATCGTGGTGTCACAAGACATCGATGGACTGGTCGA<br>TATGAGGCTCATCTATGGGATAATAGTTGTAGAAGGGAAGGC<br>CAGTCTAGGAAAGGAAGACAAGTTTACTTGGGTGGATATGA<br>CAAAGAAGATAAAGCAGCAAGATCATATGATCTAGCTGCACT<br>TAAGTACTGGGGTCCTTCAACTACTACTAATTTCCCCATTACA<br>AACTACGAGAAAGAAGTAGAGGAAATGAAGCACATGACGA<br>GACAAGAGTTCGTGGCTGCCATTAGAAGGAAAAGTAGTGGA<br>TTTTCGAGAGGCGCTTCGATGTATCGAGGAGTTACAAGGCAT<br>CACCAACATGGAAGATGGCAAGCAAGGATCGGCCGAGTCGC<br>CGGAAACAAAGACCTCTACTTGGGAACTTTTAGCACTGAGG<br>AAGAAGCAGCAGAAGCTTACGATATAGCTGCAATAAAGTTTA<br>GAGGACTTAATGCAGTGACCAACTTCGAGATCAACCGGTAC<br>GACGTGAAAGCCATTCTAGAGAGTAGCACTCTTCCCATCGGA<br>GGAGGCGCAGCTAAACGGCTCAAAGAAGCTCAAGCTCTTGA<br>GTCTTCAAGGAAACGCGAGGCGGAGATGATAGCCCTTGGTT<br>CAAGTTTCCAGTACGGTGGTGGCTCGAGCACAGGCTCTGGC<br>TCCACCTCATCAAGACTTCAGCTTCAACCTTACCCTCTAAGC<br>ATTCAACAACCATTAGAGCCTTTTCTATCTCTTCAGAACAATG<br>ACATCTCTCATTACAACAACAACAATGCTCACGATTCCTCCTC<br>TTTTAATCACCATAGCTATATCCAGACACAACTTCATCTCCAC<br>CAACAGACCAACAATTACTTGCAGCAACAGTCGAGCCAGAA<br>CTCTCAGCAGCTCTACAATGCGTATCTTCATAGCAATCCGGCT<br>CTGCTTCATGGACTTGTCTCTACCTCTATCGTTGACAACAATA<br>ATAACAATGGAGGCTCTAGTGGGAGCTACAACACTGCAGCAT<br>TTCTTGGGAACCACGGTATTGGTATTGGGTCCAGCTCGACTG<br>TTGGATCGACCGAGGAGTTTCCAACCGTTAAAACAGATTACG<br>ATATGCCTTCCAGTGATGGAACCGGAGGGTATAGTGGTTGGA<br>CCAGTGAGTCTGTTCAGGGGTCAAACCCTGGTGGTGTTTTCA<br>CTATGTGGAATGAGTAAACAAGGATCTCTTTCTTGCGGCACA<br>AGGAATGGGTCTGAATAGTAACTATTTTAATCTCTACTCTCAA<br>TTTTTATTTCTTTCTTGTCTTGGAAGTAATAGGATCAATTTAGA<br>GGGGGCCTTTTGGTAGAAGAATTTAGGTGACGCAAGGAGTT<br>CTTTCTATATTTACGGCGAAGGTATTCTCAGACTGGTTAAATG<br>GTTCTTGAGGGTTAGGGTTTTATGGGTACTCAATTAAGTTTGT<br>GTACCCAAATTTTGTTTACAGCTAACCCTTTTGACTAATTTCC<br>AAATTTGTAATTCGTTTATAAAGATGTTTTCTCTAGCTCTT |
| Cap protein sequence of TRV | 612 | ATGGGAGATATGTACGATGAATCATTTGACAAGTCGGGCGGT<br>CCTGCTGACTTGATGGACGATTCTTGGGTGGAATCAGTTTCG<br>TGGAAAGATCTGTTGAAGAAGTTACACAGCATAAAATTTGCA<br>CTACAGTCTGGTAGAGATGAGATCACTGGGTTACTAGCGGCA<br>CTGAATAGACAGTGTCCTTATTCACCATATGAGCAGTTTCCAG<br>ATAAGAAGGTGTATTTCCTTTTAGACTCACGGGCTAACAGTG<br>CTCTTGGTGTGATTCAGAACGCTTCAGCGTTCAAGAGACGA<br>GCTGATGAGAAGAATGCAGTGGCGGGTGTTACAAATATTCCT<br>GCGAATCCAAACACAACGGTTACGACGAACCAAGGGAGTAC |

| | | |
|---|---|---|
| | | TACTACTACCAAGGCGAACACTGGCTCGACTTTGGAAGAAG<br>ACTTGTACACTTATTACAAATTCGATGATGCCTCTACAGCTTT<br>CCACAAATCTCTAACTTCGTTAGAGAACATGGAGTTGAAGAG<br>TTATTACCGAAGGAACTTTGAGAAAGTATTCGGGATTAAGTT<br>TGGTGGAGCAGCTGCTAGTTCATCTGCACCGCCTCCAGCGAG<br>TGGAGGTCCGATACGTCCTAATCCC |
| Anti-CRISPR protein AcrIIA5 sequence delivered by TRV | 420 | ATGGCCTATGGCAAGTCTCGTTATAATTCCTATAGGAAACGTA<br>GCTTCAACCGATCTAACAAGCAACGTAGAGAATACGCACAG<br>GAGATGGATCGACTTGAGAAAGCATTCGAAAACCTGGACGG<br>GTGGTACCTTTCCTCAATGAAGGATAGCGCCTACAAGGATTT<br>TGGTAAGTACGAAATCCGATTGTCTAACCACTCAGCCGACAA<br>TAAATATCATGATCTCGAAAATGGGCGATTGATCGTGAACATC<br>AAGGCTTCAAAACTCAATTTTGTCGATATTATCGAAAATAAAC<br>TTGATAAGATCATCGAGAAGATCGACAAGCTCGACCTGGATA<br>AGTATAGATTTATTAACGCAACCAATCTCGAACATGACATTAA<br>GTGCTACTACAAAGGCTTCAAGACGAAAAAAGAGGTGATT |

**Supplementary Table 4. Guide RNA sequences used in this study.**

| Name | Length (nt) | Sequence |
|---|---|---|
| gRNA_01 | 41 | GGAGGAGTCCTGGGTAACGGTAACAACACCACCGTCTTCGA |
| gRNA_02 | 41 | TGGTACCACGCAGTTTAACTTTGTAGATGAACTCACCGTCT |
| gRNA_03 | 82 | GGAGGAGTCCTGGGTAACGGTAACAACACCACCGTCTTCGA<br>AGTTCATAACACGTTCCCATTTGAAACCTTCCGGGAAGGAC |
| gRNA_04 | 82 | CATGGTTTTTTCTGCATAACCGGACCGTCGGACGGGAAGTT<br>GGTACCACGCAGTTTAACTTTGTAGATGAACTCACCGTCT |
| gRNA_05 | 82 | CCGGTTTGCTCACACCGAGAAAATCGGCCACTTTTGGAACCT<br>CTCCTCCTTCATCTCCTCCTCCACCGTGTGGATTGATCAT |
| gRNA_06 | 82 | TTGATTGGACGCTAGGCATCAAGCTATTGGTATGGAAGTAGT<br>AGTCTGAGTCGTTGTAAGCTACTAGGTGGTTGGATTGGTT |
| gRNA_07 | 123 | GGAGGAGTCCTGGGTAACGGTAACAACACCACCGTCTTCGA<br>AGTTCATAACACGTTCCCATTTGAAACCTTCCGGGAAGGACA<br>GTTTCAGGTAGTCCGGGATGTCAGCCGGGTGTTTAACGTA |
| gRNA_08 | 123 | CCGTCTTCCGGGTACATACGTTCGGTGGAAGCTTCCCAACCC<br>ATGGTTTTTTCTGCATAACCGGACCGTCGGACGGGAAGTTG<br>GTACCACGCAGTTTAACTTTGTAGATGAACTCACCGTCT |
| gRNA_09 | 164 | GGAGGAGTCCTGGGTAACGGTAACAACACCACCGTCTTCGA<br>AGTTCATAACACGTTCCCATTTGAAACCTTCCGGGAAGGACA<br>GTTTCAGGTAGTCCGGGATGTCAGCCGGGTGTTTAACGTAAG<br>CTTTGGAACCGTACTGGAACTGCGGGGACAGGATGTCCC |
| gRNA_10 | 164 | CTTTCAGTTTCAGACGCATTTTGATTTCACCTTTCAGAGCACC<br>GTCTTCCGGGTACATACGTTCGGTGGAAGCTTCCCAACCCAT<br>GGTTTTTTCTGCATAACCGGACCGTCGGACGGGAAGTTGGT<br>ACCACGCAGTTTAACTTTGTAGATGAACTCACCGTCT |
| gRNA_11 | 325 | GGAGGAGTCCTGGGTAACGGTAACAACACCACCGTCTTCGA<br>AGTTCATAACACGTTCCCATTTGAAACCTTCCGGGAAGGACA<br>GTTTCAGGTAGTCCGGGATGTCAGCCGGGTGTTTAACGTAAG<br>CTTTGGAACCGTACTGGAACTGCGGGGACAGGATGTCCCAA<br>GCGAACGGCAGCGGACCACCTTTGGTAACTTTCAGTTTAGC<br>GGTCTGGGTACCTTCGTACGGACGACCTTCACCTTCACCTTC |

| | | |
|---|---|---|
| | | GATTTCGAACTCGTGACCGTTAACGGAACCTTCCATACGAACTTTGAAACGCATGAACTCTTTGATAACGTCTTCG |
| gRNA_12 | 325 | GGAGTGACGACCTTCAGCACGTTCGTACTGTTCAACGATGGTGTAGTCTTCGTTGTGGGAGGTGATGTCCAGTTTGATGTCGGTTTTGTAAGCACCCGGCAGCTGAACCGGTTTTTAGCCATGTAGGTGGTTTTAACTTCAGCGTCGTAGTGACCACCGTCTTTCAGTTTCAGACGCATTTTGATTTCACCTTTCAGAGCACCGTCTTCCGGGTACATACGTTCGGTGGAAGCTTCCCAACCCATGGTTTTTTTCTGCATAACCGGACCGTCGGACGGGAAGTTGGTACCACGCAGTTTAACTTTGTAGATGAACTCACCGTCT |
| gRNA_13 | 82 | AAACTGATTCCACCCAAGAATCGTCCATCAAGTCAGCAGGACCGCCCGACTTGTCAAATGATTCATCGTACATATCTCCCAT |
| gRNA_14 | 82 | GTGCCGCTAGTAACCCAGTGATCTCATCTCTACCAGACTGTAGTGCAAATTTTATGCTGTGTAACTTCTTCAACAGATCTTT |
| gRNA_15 | 82 | GTGCGTATTCTCTACGTTGCTTGTTAGATCGGTTGAAGCTACGTTTCCTATAGGAATTATAACGAGACTTGCCATAGGCCAT |
| gRNA_16 | 82 | CAAAATCCTTGTAGGCGCTATCCTTCATTGAGGAAAGGTACCACCCGTCCAGGTTTTCGAATGCTTTCTCAAGTCGATCCAT |

**Supplementary Table 5. Sequences of other biological components used in this study.**

| Components | Sequence (5' to 3') |
|---|---|
| sfGFP | ATGAGCAAAGGAGAAGAACTTTTCACTGGAGTTGTCCCAATTCTTGTTGAATTAGATGGTGATGTTAATGGGCACAAATTTTCTGTCCGTGGAGAGGGTGAAGGTGATGCTACAAACGGAAAACTCACCCTTAAATTTATTTGCACTACTGGAAAACTACCTGTTCCGTGGCCAACACTTGTCACTACTCTGACCTATGGTGTTCAATGCTTTTCCCGTTATCCGGATCACATGAAACGGCATGACTTTTTCAAGAGTGCCATGCCCGAAGGTTATGTACAGGAACGCACTATATCTTTCAAAGATGACGGGACCTACAAGACGCGTGCTGAAGTCAAGTTTGAAGGTGATACCCTTGTTAATCGTATCGAGTTAAAGGGTATTGATTTTAAAGAAGATGGAAACATTCTTGGACACAAACTCGAGTACAACTTTAACTCACACAATGTATACATCACGGCAGACAAACAAAAGAATGGAATCAAAGCTAACTTCAAAATTCGCCACAACGTTGAAGATGGTTCCGTTCAACTAGCAGACCATTATCAACAAAATACTCCAATTGGCGATGGCCCTGTCCTTTTACCAGACAACCATTACCTGTCGACACAATCTGTCCTTTCGAAAGATCCCAACGAAAAGCGTGACCACATGGTCCTTCTTGAGTTTGTAACTGCTGCTGGGATTACACATGGCATGGATGAGCTCTACAAATAA |
| eYGFPuv | ATGACAACCTTCAAAATCGAGTCCCGGATCCACGGCAACCTCAACGGGGAGAAGTTCGAGTTGGTTGGAGGTGGAGTAGGTGAGGAGGGTCGCCTCGAGATTGAGATGAAGACTAAAGATAAACCACTGGCATTCTCTCCCTTCCTGCTGACCACTTGCATGGGTTACGGGTTCTACCACTTCGCCAGCTTCCCAAAGGGGATTAAGAACATCTATCTTCATGCTGCAACGAACGGAGGTTACACCAACACCAGGAAGGAGATCTATGAAGACGGCGGCATCTTGGAGGTCAACTTCCGTTACACTTACGAGTTCAACAAGATCATCGGTGACGTCGAGTGCATTGGACATGGATTCCCAAGTCAGAGTCCGATCTTCAAGGACACGATCGTGAAGAGTTGTCCCACG |

| | |
|---|---|
| | GTGGACCTGATGTTGCCAATGTCCGGGAACATCATCGCCAG CTCCTACGCTTACGCCTTCCAACTGAAGGACGGCTCTTTCTA CACGGCAGAAGTCAAGAACAACATAGACTTCAAGAATCCA ATCCACGAGTCCTTCTCGAAGTCAGGGCCCATGTTCACCCA CAGACGTGTCGAGGAGACTCTCACCAAGGAGAACCTTGCC ATAGTGGAGTACCAGCAGGTTTTCAACAGCGCCCCAAGAGA CATGTAG |
| P2A | GCTACTAACTTCAGCCTGCTGAAGCAGGCTGGAGACGTGGA GGAGAACCCTGGACCT |
| GSG linker | GGAAGCGGA |
| HP14 | ACGTCGACTCTCGAGTGAGATTGTTGACGGTACCGTATTTT |
| IGS1 | GGGTCA |
| IGS2 | GTAGAT |
| P1_loop | AAATAGCAATATTTACCTTT |
| P1_loop_01 | AAATAGCAA |
| P1_loop_02 | TAGTTACCTTT |
| Stem_01 | GGATCA |
| Stem_02 | TGATCC |
| Ribozyme | AAAAGTTATCAGGCATGCACCTGGTAGCTAGTCTTTAAACC AATAGATTGCATCGGTTTAAAAGGCAAGACCGTCAAATTGC GGGAAAGGGGTCAACAGCCGTTCAGTACCAAGTCTCAGGG GAAACTTTGAGATGGCCTTGCAAAGGGTATGGTAATAAGCT GACGGACATGGTCCTAACCACGCAGCCAAGTCCTAAGTCAA CAGATCTTCTGTTGATATGGATGCAGTTCACAGACTAAATGT CGGTCGGGGAAGATGTATTCTTCTCATAAGATATAGTCGGAC CTCTCCTTAATGGGAGCTAGCGGATGAAGTGATGCAACACT GGAGCCGCTGGGAACTAATTTGTATGCGAAAGTATATTGATT AGTTTTGGAGTACTCG |
| dRibozyme | AAAAGTTATCAGGCATGCACCTGGTAGCTAGTCTTTAAACC AATAGATTGCATCGGTTTAAAAGGCAAGACCGTCAAATTGC GGGAAAGGGGTCAACAGCCGTTCAGTACCAAGTCTCAGGG GAAACTTTGAGATGGCCTTGCAAAGGGTATGGTAATAAGCT GACGGACATGGTCCTAACCACGCAGCCAAGTCCTAAGTCAA CAGATCTTCTGTTGATATGGATGCAGTTCACAAACTAAATGT CGGTCGGGGAAGATGTATTCTTCTCATAAGATATAGTCGGAC CTCTCCTTAATGGGAGCTAGCGGATGAAGTGATGCAACACT GGAGCCGCTGGGAACTAATTTGTATGCGAAAGTATATTGATT AGTTTTGGAGTACTCG |
