## Supplementary figures for "Ribozyme-based biosensor for imaging gene expression in plants"

### Extended Data Fig. 1

(**a**) Bright field of pictures in Fig. 1d. (**b**) Bright field of pictures in Fig. 1e. The pictures were taken with a Zeiss LSM 710 confocal microscope under bright light. Scale bar = 100 μm.

a

35s::HP14::sfGFP 35s::HP14::R_sfGFP 35s::HP14::dR_sfGFP

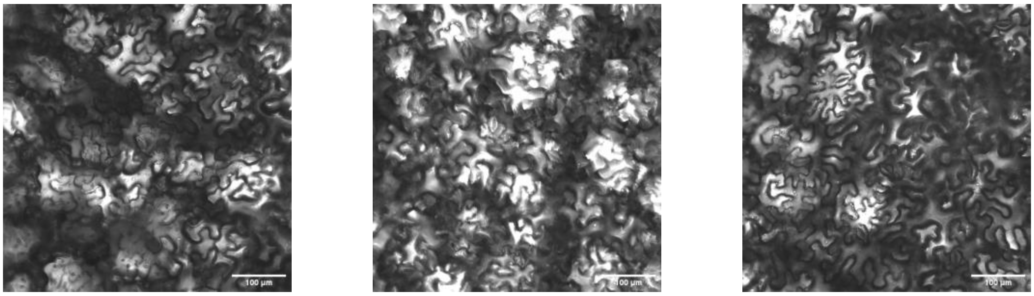

b

35s::sfGFP 35s::R_sfGFP 35s::dR_sfGFP

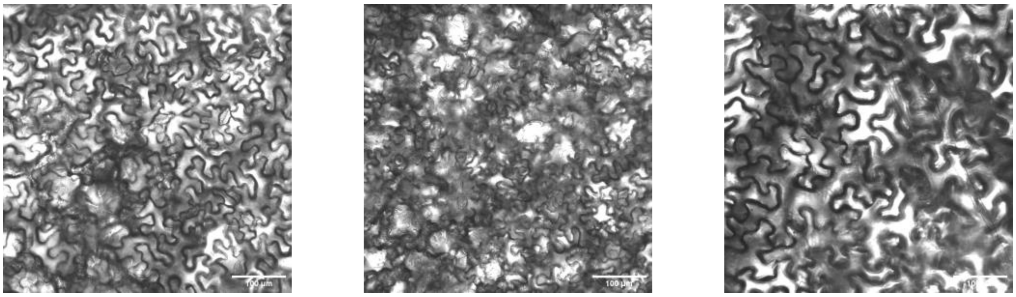

### Extended Data Fig. 2

(**a**) Bright field of pictures in Fig. 2c. (**b**) Florescent signals and bright field of split-ribozyme system targeting RFP transcript with different gRNA length. The pictures were taken with a Zeiss LSM 710 confocal microscope. Scale bar = 100 µm.

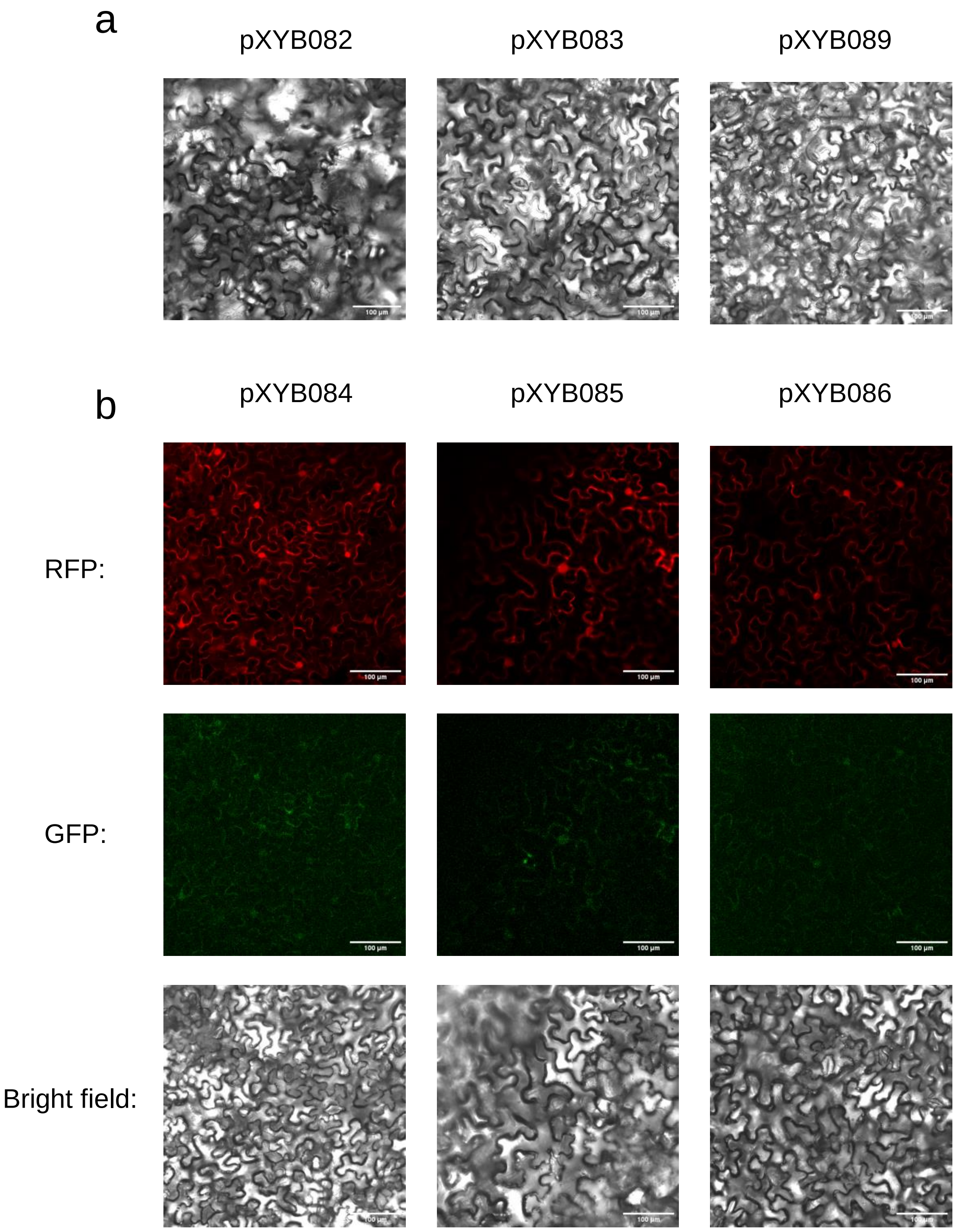

### Extended Data Fig. 3

(**a**) Bright field of pictures in Fig. 3b. (**b**) Bright field of pictures in Fig. 3c. The pictures were taken with a Zeiss LSM 710 confocal microscope under bright light. Scale bar = 100 μm.

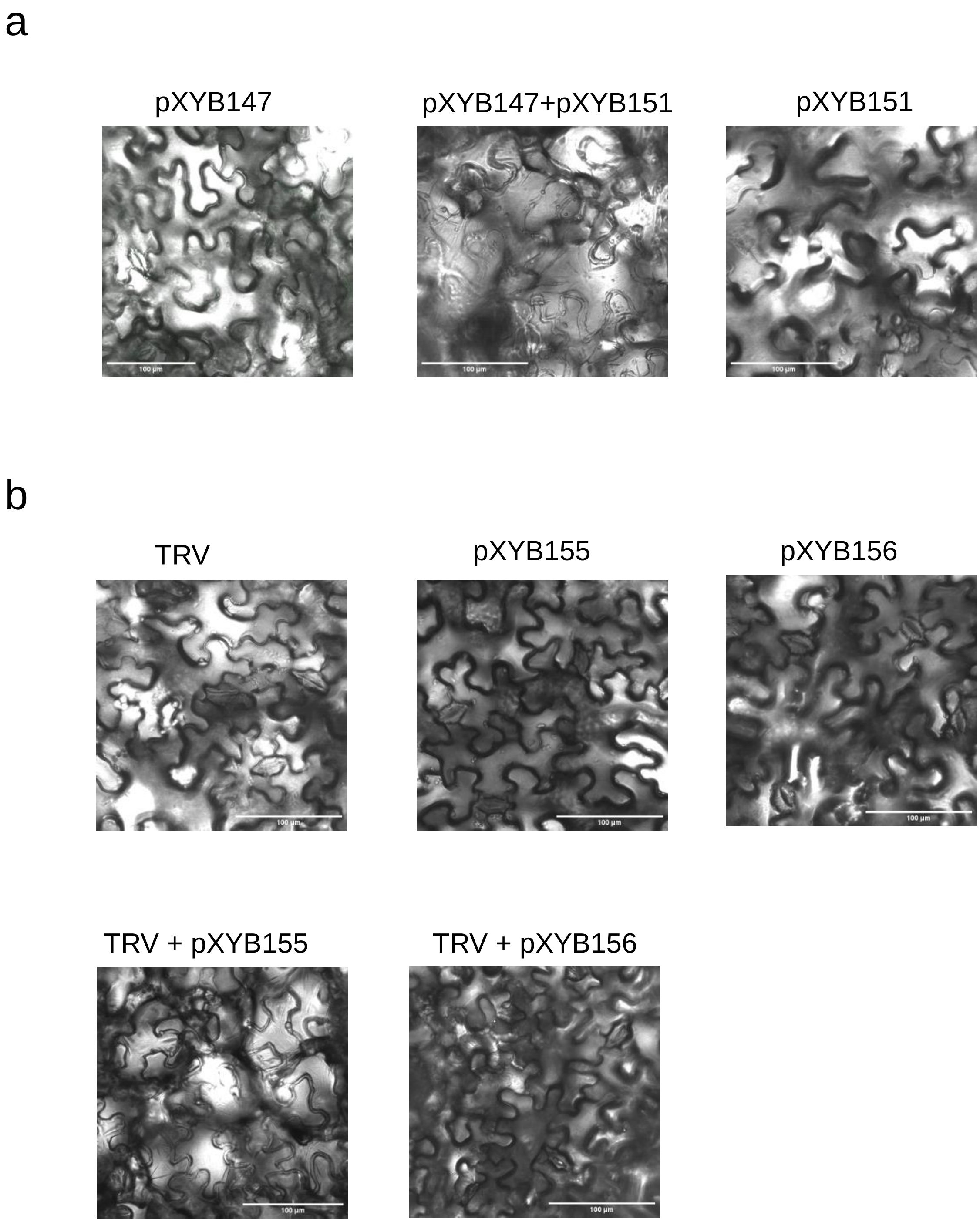

### Extended Data Fig. 4

Green fluorescent signal observed in the roots of representative transgenic *Arabidopsis thaliana* plants. The pictures were taken with a Zeiss LSM 710 confocal microscope. Scale bar = 100 μm.

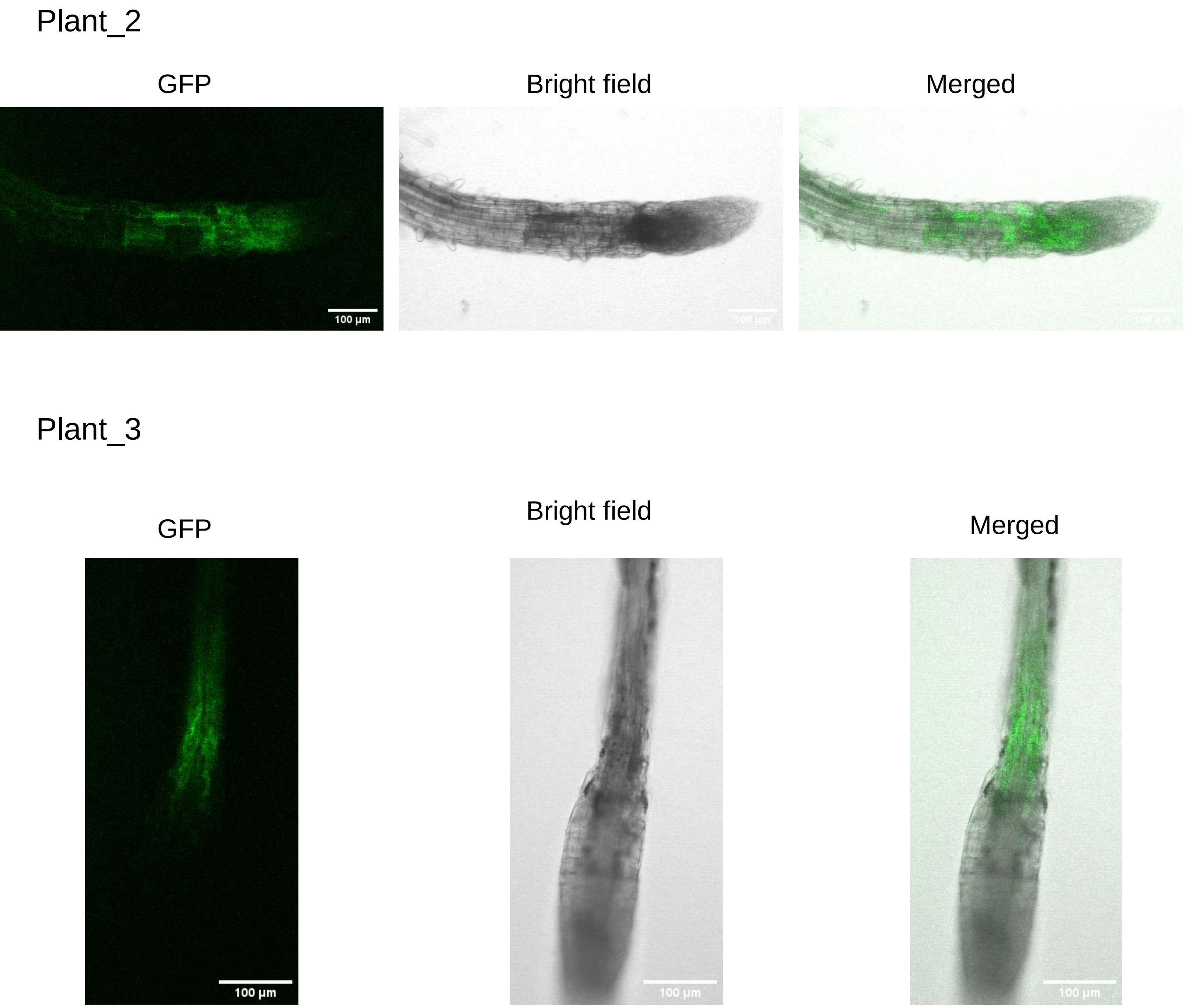

### Extended Data Fig. 5

(a) Construct design for testing different RNA stability motifs and new design of Ribozyme with new internal guide sequence (IGS). 35S: Promoter region of the Cauliflower mosaic virus *35S* gene; 5’UTR: 5′ untranslated region of *Arabidopsis thaliana COLD REGULATED 47* gene; HSP: terminator region of *A. thaliana HEAT SHOCK PROTEIN 18.2* gene; GFPuv: enhanced *YELLOW GREEN FLUORESCENT LIKE PROTEIN*; R with black rectangle: ribozyme; dR with dark red rectangle: a catalytically dead G264A mutant ribozyme with IGS and PI loop removed; Blue rectangle: tEvopreq motif; Pink rectangle: gq2 motif; Brown rectangle: Triplex motif; R with yellow rectangle: Ribozyme with IGS2; dR with green rectangle: a catalytically dead G264A mutant ribozyme with IGS2. (b) Visualization of eYGFPuv in leaf under UV light, three days post *N. benthamiana* leaf infiltration.

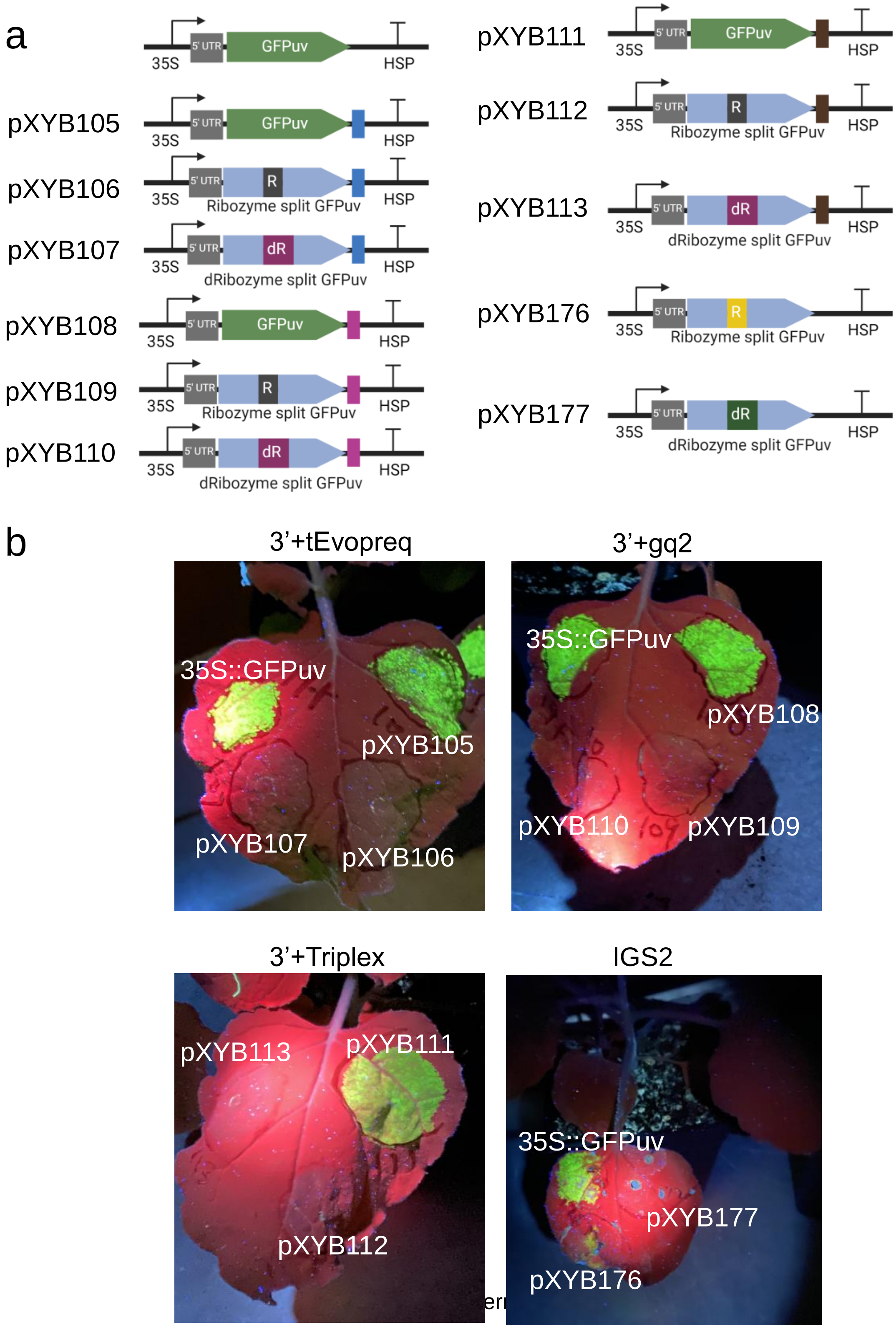

### Extended Data Fig. 6

Bright field of pictures in Fig. 4d. The pictures were taken with a Zeiss LSM 710 confocal microscope under bright light. Scale bar = 100 µm.

pXYB164

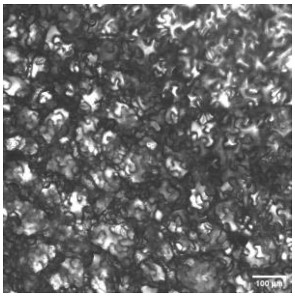

pXYB162

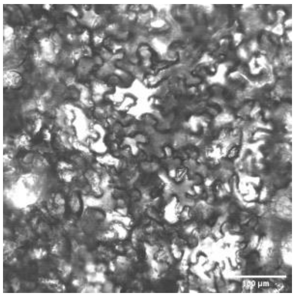

pXYB163

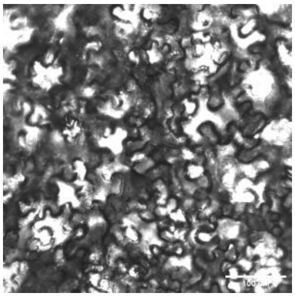

### Extended Data Fig. 7

Illustration of the potential applications of the “Plant RNA Vision” technology. Through various delivery methods, the “Plant RNA Vision” toolkit can be used for detection of gene expression at cellular, tissue/organ, and whole-plant levels during plant growth, development, and response to abiotic/biotic factors.

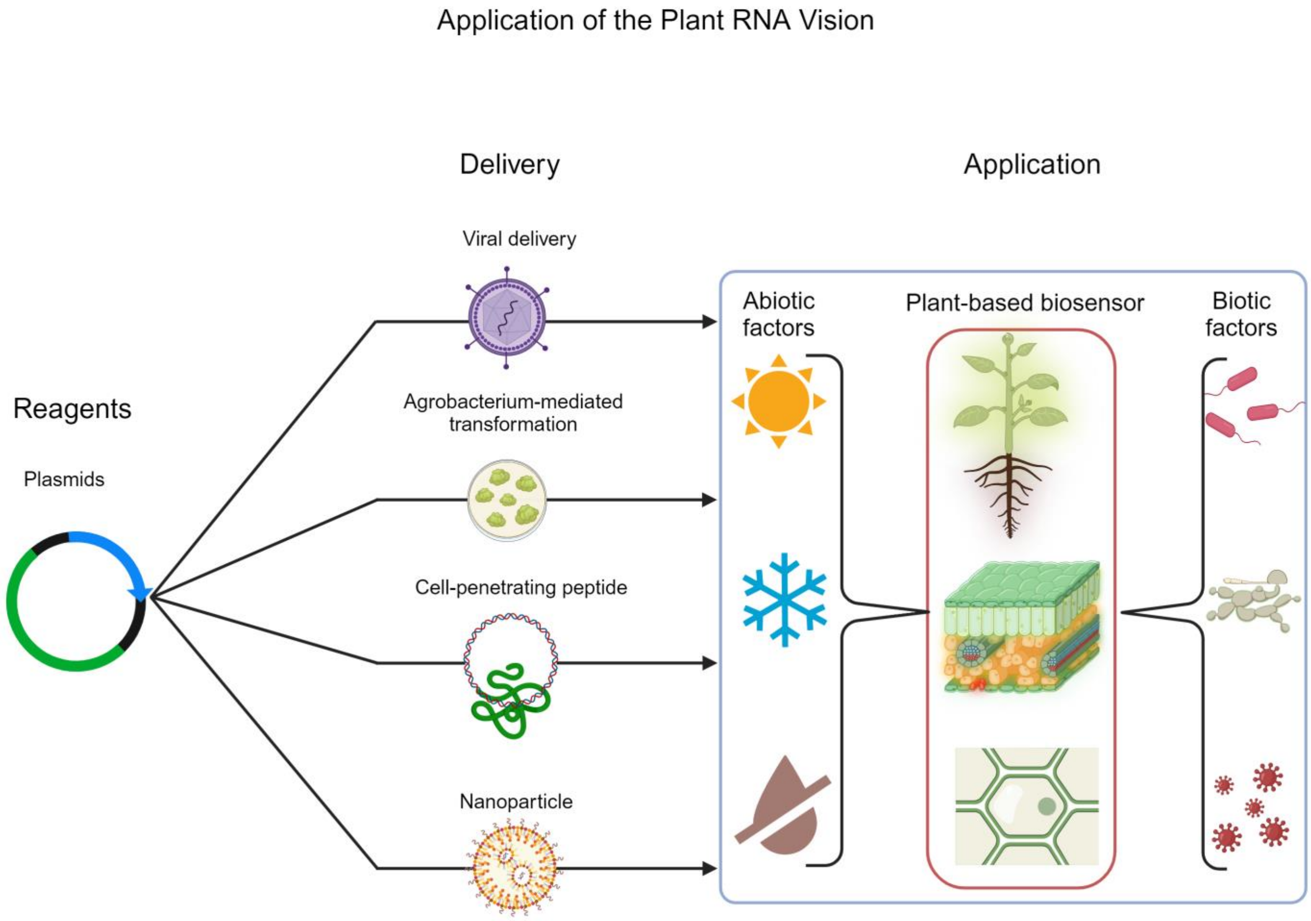

### Extended Data Fig. 8

Illustration of guide RNA design.

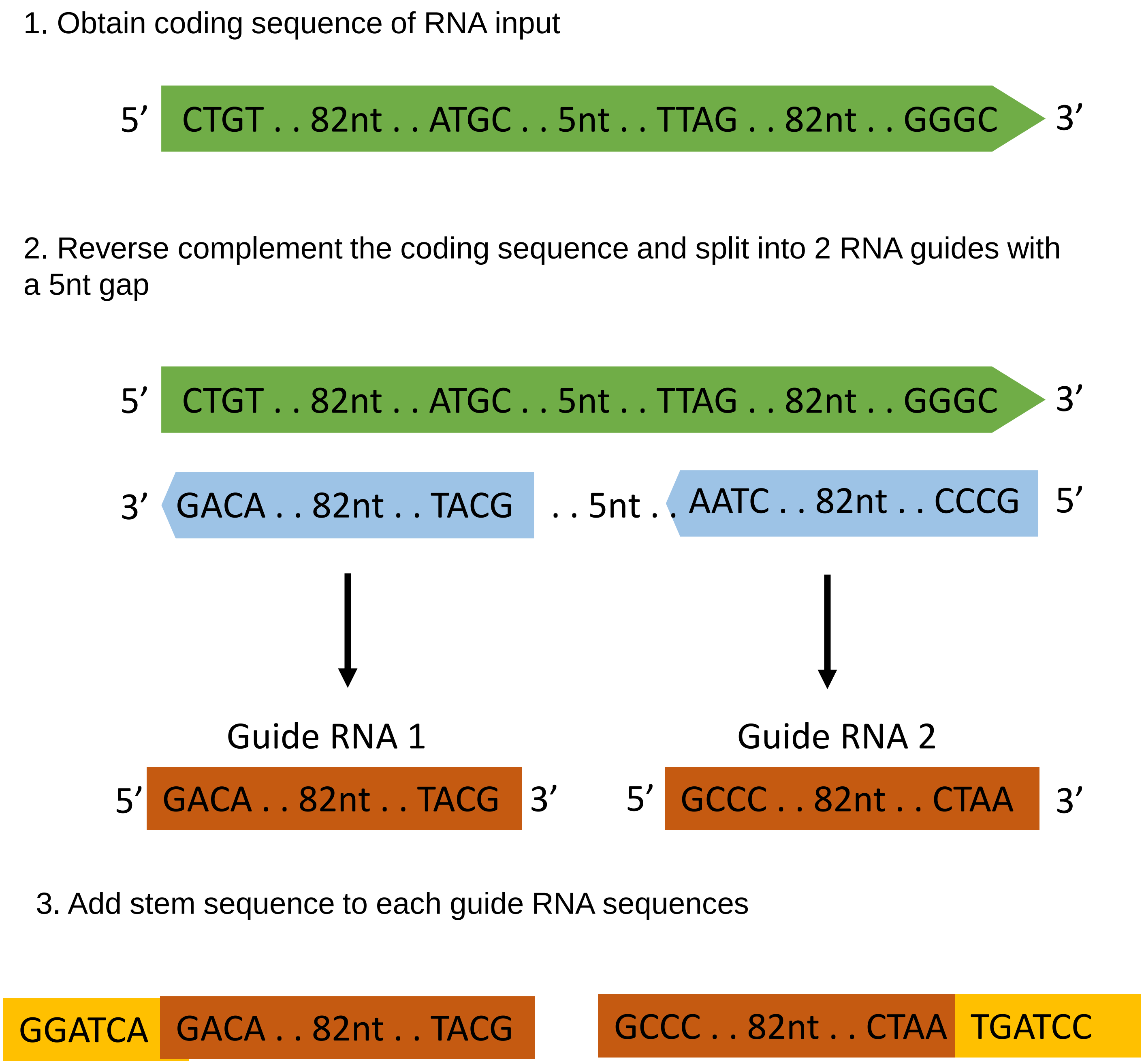


---Internal Use---
